# Movement-assisted localization from acoustic telemetry data

**DOI:** 10.1101/2019.12.31.890962

**Authors:** Nathan J. Hostetter, J. Andrew Royle

**Affiliations:** U.S. Geological Survey, Patuxent Wildlife Research Center, Laurel, MD 20708; Washington Cooperative Fish and Wildlife Research Unit, School of Aquatic and Fishery Sciences, University of Washington, Seattle, WA, 98195

**Keywords:** Acoustic telemetry, Bioacoustics, Movement ecology, Sound attenuation, Spatial capture-recapture, Telemetry

## Abstract

**Background:** Acoustic telemetry technologies are being rapidly deployed to study a variety of aquatic taxa including fishes, reptiles, and marine mammals. Large cooperative telemetry networks produce vast quantities of data useful in the study of movement, resource selection and species distribution. Efficient use of acoustic telemetry data requires estimation of acoustic source locations from detections at sensors (i.e. localization). Multiple processes provide information for localization estimation including detection/non-detection data at sensors, information on signal rate, and an underlying movement model describing how individuals move and utilize space. Frequently, however, localization methods only integrate a subset of these processes and do not utilize the full spatial encounter history information available from sensor arrays.

**Methods:** In this paper we draw analogies between the challenges of acoustic telemetry localization and newly developed methods of spatial capture-recapture (SCR). We develop a framework for localization that integrates explicit sub-models for movement, signal (or cue) rate, and detection probability, based on acoustic telemetry spatial encounter history data. This method, which we call movement-assisted localization, makes efficient use of the full encounter history data available from acoustic sensor arrays, provides localizations with fewer than three detections, and even allows for predictions to be made of the position of an individual when it was *not* detected at all. We demonstrate these concepts by developing generalizable Bayesian formulations of the SCR movement-assisted localization model to address study-specific challenges common in acoustic telemetry studies.

**Results:** Simulation studies show that movement-assisted localization models improve point-wise RMSE of localization estimates by > 50% and greatly increased the precision of estimated trajectories compared to localization using only the detection history of a given signal. Additionally, integrating a signal rate sub-model reduced biases in the estimation of movement, signal rate, and detection parameters observed in independent localization models.

**Conclusions:** Movement-assisted localization provides a flexible framework to maximize the use of acoustic telemetry data. Conceptualizing localization within an SCR framework allows extensions to a variety of data collection protocols, improves the efficiency of studies interested in movement, resource selection, and space-use, and provides a unifying framework for modeling acoustic data.

## 1 Background

The use of acoustic telemetry has expanded rapidly in recent years. Acoustic telemetry has been widely adopted in studies of marine mammals, reptiles (e.g., sea turtles, crocodiles), and fish and has become an important tool in the study of spatial ecology in marine and freshwater systems [1–3]. To address these challenges large-scale cooperative telemetry networks are now deployed around the world [4–6] including the Great Lakes Acoustic Telemetry Observation System [7], the Atlantic Cooperative Telemetry Network [8], and the Ocean Tracking Network [9].

A key objective of acoustic telemetry studies is *localization* of sources using data from arrays of receivers, i.e., estimation of the location of an individual source from detections at one or more sensors of an array. When regarded formally as an estimation problem, localization is essentially statistical triangulation, and it can be done when signals are obtained from an array of sensors configured so that multiple detections of the same signal are possible [10–15]. Localization may be based on simple detection history information (the pattern of sensors at which detections occur) and also auxiliary information on time delay of arrival at different sensors, or signal strength. Localization of sources is crucial for studies of spatial ecology, resource selection, and density estimation. The precision of localization is therefore an important objective function in many acoustic studies, where researchers often consider trade-offs in the number and spacing of sensors to optimize precision of localizations with logistical and financial constraints [16].

Multiple types of data can provide information important to localization from acoustic arrays. For example, detections of signals at fixed sensors provide information on individual source locations. Similarly, non-detections also provide information on the location of an individual, however, most localization approaches do not use observed zeros – that is, locations of sensors where detections did not occur, and in some cases do not use occasions when individuals are detected at fewer than 3 sensors. Second, the location of an individual at time *t* should be informed to some extent by data from previous and subsequent times, with the degree of information decreasing as the time interval between observed source locations increases. Most methods of localization, however, do not explicitly integrate information about movement processes to inform the localization of acoustic sources [but see 17].

Integrating movement processes and spatial capture-recapture [SCR, 18] data in terrestrial systems has led to important methodological improvements that are relevant to localization in acoustic telemetry [e.g., 19–21]. Indeed, localization is analogous to inference about the activity or home-range center in SCR models, and therefore SCR ideas have been adapted to accommodate data obtained by acoustic sampling methods [15, 19, 22, 23]. The benefit of this SCR-based view of localization, what we refer to as statistical localization, is that it allows *in situ* estimation of parameters related to detection range along with simultaneous localization of source locations, and potentially other parameters that describe the detection process, the rate at which signals are produced, the distribution of individuals, and their movement through time [21, 24–26]. Moreover, SCR-based statistical localization uses all available detection history information, including the observed non-detections, and sources which are detected by only 1 or 2 sensors. Herein, we propose to modify and extend these SCR ideas to describe a general conceptual framework to localization in acoustic telemetry systems which integrates detection data from sensor arrays with explicit models of individual movement and signal (or “cue”) rates. The important advance of our work is recognition that integrating an explicit movement model with the estimation of source locations from spatial encounter histories introduces additional information into the localization process. In general, we believe that integrating an explicit movement model to link locations through time will improve localization, provide deeper insight into movement dynamics, resource selection and individual behavior, and allow coarser sensor spacing which may improve sensor array design. Moreover, formulating the localization process in terms of a spatially explicit model of individual distribution, movement, and signal rates may lead to solutions to some similar outstanding problems in applications of acoustic monitoring related to estimation of density, resource selection, and movement.

## 2 Methods

### 2.1 Data structure and model

Let *u_t_* be the unknown location at the time the *t^th^* signal was produced. At times *t* = 1,…,*T* the transmitter produces a signal which may be detected by one or more sensors in an array. Let Δ_*t*_ be the interval between signal transmissions (hereafter “signals” or “transmissions”). In some cases Δ_*t*_ is constant for all tags and prescribed by design but, in practice, when many tags are deployed at the same frequency the interval is often set to be random in order to introduce an offset in detection times of individuals and avoid interference among tags. For example, an individual tag might be set to emit a signal on a random schedule with a minimum of 50 seconds and a maximum of 100 seconds between signals. Thus Δ_*t*_ ~ Uniform(50, 100).

For demonstration purposes, we assume a Markovian movement process conditional on Δ_*t*_ according to (e.g., Brownian motion)

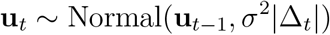

although we note that a number of alternative movement models are possible [27]. The observed data for each time *t* are the locations of the sensors that detected the individual (including possibly none). To be consistent with SCR terminology we call this the spatial encounter history and denote it by the vector **y**_*t*_ where elements *y_j,t_* = 1 if the individual tag was detected by receiver *j* at transmission *t*. Receivers have coordinates **x**_*j*_ which are fixed by design. In some cases we might have time-difference-of-arrival (TDOA) information but, in practice, such information may not be available and, instead, only a coarse summary of arrival time is given [e.g., rounded to whole second, 15]

### 2.2 Localization based on the observed encounter history

Using only the spatial encounter history information, localization is achieved by application of Bayes’ rule to compute the conditional probability of **u**_*t*_ given the observed encounter history **y**_*t*_, i.e., the posterior probability distribution of the latent source location **u**_*t*_,

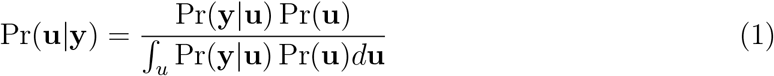

This requires specification of two probability distributions: (1) Pr(**u**) is the probability distribution of the source location which, lacking specific additional information, can be taken to be uniform over the planar region in the vicinity of the acoustic array^1^. If explicit habitat features are available then parameters which allow for non-uniformity can be estimated [28, 29]. (2) We also require the probability distribution of the spatial encounter history

Pr(**y**|**u**) which, for binary detection encounter data, is determined by a set of detection probabilities *p_j,t_*, normally taken to be a homogeneous function of distance between the source location **u** and the sensor locations **x**_*j*_. For example, the normal kernel is commonly used in distance sampling [30] and spatial capture-recapture applications:

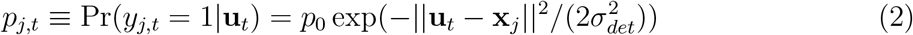

which has parameters *p*_0_ and 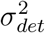. In acoustic telemetry applications a logistic model is often used [31]

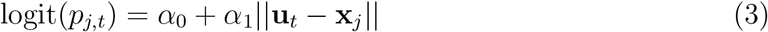

The parameters of either encounter model can be estimated by maximum likelihood without difficulty [19, 32] and used as a plug-in or empirical Best Unbiased Predictor (BUP) [33] of the latent variable **u**_*t*_ based on the posterior distribution (Eq. 1). From a practical standpoint the functional form of the detection model is not important, but note that these standard models are a function of some power of Euclidean distance.

In practice, formal statistical estimation of the parameters from observed acoustic data is seldom done. Instead, the parameters are prescribed based on estimates from controlled “range testing” studies [e.g., 31] in which transmitters are located at prescribed distances from one or more sensors and the detection of broadcast signals is modeled given known source locations. However, we believe that a better approach is to use formal models of the observed encounter history data in a spatial capture-recapture framework and estimate detection parameters directly from the observed data [e.g., following 19, 20].

### 2.3 Movement-assisted localization

An obvious shortcoming of the approach described above is that localization of **u**_*t*_ only uses information at time *t*. Here, occasions with only 1 or 2 detections are sometimes not localized due to insufficient data, or alternatively, detection data are binned into longer time windows until a sufficient number of detections are available for localization. In the latter case the localization is producing an estimate of average location during the full time window. The obvious trade-off from a design standpoint is more detections at *t* improves localization, which is often accomplished by placing sensors closer together, thus requiring more sensors to sample a given area and greater cost. On the other hand, increasing the sensor spacing produces fewer detections, and fewer and more imprecise localizations.

We propose to resolve this trade-off formally by extending the localization model described above through the integration of an explicit movement model to simultaneously estimate the movement and detection processes. For example, instead of assuming Pr(**u**_*t*_) is uniform we replace this assumption with the Markovian movement assumption given above, that is

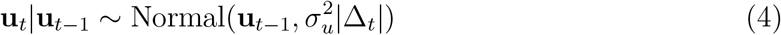

Then, in effect, the data from the previous (and subsequent) interval provide information about **u**_*t*_ via the prior distribution Pr(**u**_*t*_|**u**_*t*−1_).

Use of this movement model based prior distribution is effectively an informative prior in the sense that it restricts potential states of **u**_*t*_ to be in the vicinity of the previous state where the extent of this vicinity is determined by the parameter 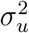 as well as the sampling interval Δ_*t*_. Thus, note that as Δ_*t*_ increases, the information provided by previous states diminishes rapidly and the prior tends to a uniform (non-informative) prior defined by the state-space.

When the signal schedule is known (see below), so that the non-detections are in effect observed, then the observed data are the spatial encounter histories **y**_*t*_ for a given individual, for each *t* = 1, 2, …, *T*. This may include all zero observations (**y**_*t*_ is a vector of zeros, indicating non-detection). Then, the joint distribution of the observed data and the latent movement tra jectory **u**_1_, … , **u**_*T*_ is

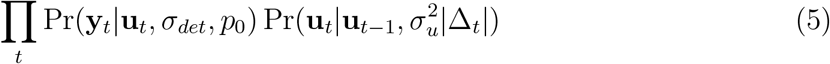

The inference objective is to jointly estimate the model parameters *p*_0_, *σ_det_*, and *σ_u_* as well as the latent trajectory **u**_1_, …, **u**_*T*_. For this we adopt a Bayesian approach based on Markov chain Monte Carlo (MCMC) as described in Section 2.3.1. When multiple tags are in operation simultaneously, data from all tags can be pooled for joint estimation of the parameters and latent trajectories from all encounter history data. In this case the encounter data are **y**_*i,t*_ and the trajectories are **u**_*i,t*_ for individual *i* and *t^th^* signal. Then the joint distribution to be analyzed is:

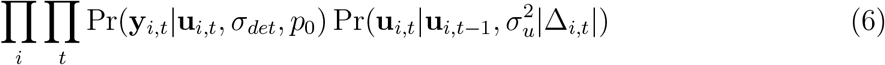

#### 2.3.1 Inference for the movement-assisted localization model

In general, it is challenging to express the likelihood of the observed data (the spatial encounter histories including auxiliary information such as arrival time) as a function of the model parameters, because the model is formulated explicitly in terms of latent (unobserved) locations, the ut variables. The problem has many analogs in the statistical literature, as a Hidden Markov model [34], which suggests promise toward achieving a solution to constructing the likelihood. Instead, we suggest that Bayesian analysis of the model using standard methods of Markov chain Monte Carlo (MCMC) is relatively straightforward. This is facilitated by the use of existing software packages such as JAGS [35] which requires little more than a pseudo-code representation of the model. The model described in the previous section, where the interval duration schedule is known, is shown in Panel 1. This illustrates the simplicity of the movement-assisted localization framework which amounts to formal integration of a movement model with a spatial encounter model.

Here, we seek to estimate the model parameters, which include the detection range parameter *σ_det_*, the detection probability parameter *p*_0_ and also the movement variance parameter 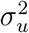. However, the key quantities we wish to estimate in the context of localization are the posterior distributions for the locations of the tag at *each* signal time *t* = 1, 2,…,*T* given all available data. In the context of Bayesian analysis, localization is naturally achieved using the posterior distribution obtained by MCMC sampling. A key step in the construction of a Metropolis-within-Gibbs algorithm is that the full-conditional distribution for **u**_*t*_ can be constructed by noting that,

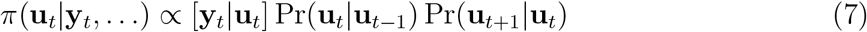

This does not have a convenient simplified form (to the best of our knowledge) but it does emphasize the point that information about the state of **u**_*t*_ derives not only from the data **y**_*t*_ but also from the previous (**u**_*t*−1_) and subsequent (**u**_*t*+1_) states. In turn, those previous and subsequent states are informed by detection data from *t* − 1 and *t* + 1, respectively. Thus our generalized localization model is using “all the data” in a manner that is prescribed by the specific movement and detection model imposed upon the system.

#### 2.3.2 Unknown duration between signals

In many acoustic telemetry applications the time interval between signals will not be known. Rather, devices are programmed to emit a signal on a random schedule, e.g., Δ_*t*_ ~ Uniform(*a, b*), but the timing and number of such signals is not registered. In this case, when the interval between *observed* signals is long, there is uncertainty in the number of missed signals. The challenge of random intervals between signals in acoustic telemetry has many similarities to the estimation of cue rates in passive acoustic studies [23]. In both instances, an individual (or an individual’s tag) is producing a cue at some partially known rate, but the number of undetected cues remains unknown. For example, if the signal schedule for a telemetry tag is random with an interval of between 90 and 150 seconds and you observe a 10 minute gap between detections of a given tag, then there are “missed signals” which have to be accommodated to achieve unbiased estimates of the detection process. Clearly it is uncertain whether the 10 minute interval contained 3, 4, 5 or 6 signals that were missed. In this case, one option for including the data obtained at the 10 minute interval is to set Δ_*t*_ = 10 and then eq 4 is correct for the observed interval. The effects of missed signals in this approach,however, is to decrease the information about the current state **u**_*t*_ provided by previous and subsequent states and biased estimates of the true detection parameters as all missed signals are ignored.

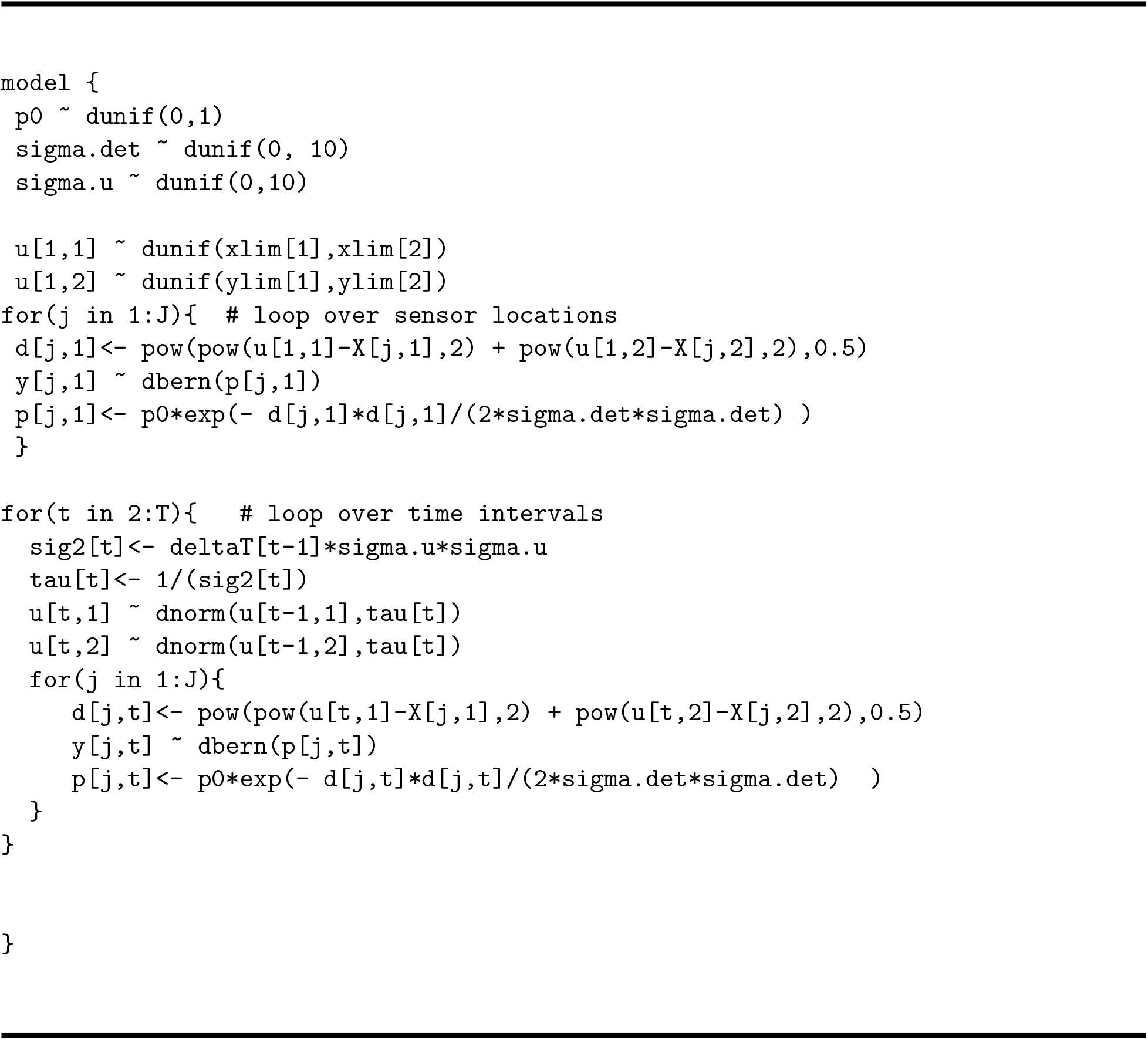

Panel 1: JAGS specification of the localization model using a Brownian motion prior distribution on animal locations. Input data are the detection history data *y*, a *J* × *T* matrix, the time interval information *T* where Δ_*t*_ is the interval between signal *t* − 1 and *t*, and the sensor locations **X**.

Alternatively, we develop an MCMC algorithm that treats the number of missed signals and the interval duration between each successive missed signal as random variables that are estimated as part of the model. To do this, we integrate an additional sub-model to describe the signal rate and associated interval durations. The specific form of the signal rate subm-odel can be tailored to any study (e.g., programmed acoustic tags versus passive cetacean cues), but generally requires two stochastic components to account for: (i) the number of missed signals and (ii) the interval length between missed signals. The R code is provided in Appendix A, while the signal rate sub-model is briefly described below.

The signal rate sub-model can be tailored to different protocols and formulated based on inter-signal rates (Δ_*t*_; e.g., defined transmitter settings in telemetry studies) or number of cues per unit time (*n*; e.g., animal calls). In most applications, assuming a distribution for interval duration also induces a distribution for the total number of missed signals and vice versa. For demonstration purposes, we develop a signal rate sub-model for acoustic tags programmed with signal intervals Δ_*t*_ ~ Uniform(*a, b*). The constraints for estimating the number of missed signals (*n*) and their associated intervals (Δ_*t*,1:*n*_) quickly becomes complex as for any observed interval, (Δ_*obs*_, i.e. the length of the *gap*), there is a known minimum and maximum number of missed signals, a fixed minimum and maximum true interval duration, and a requirement that the intervals must sum to Δ_*obs*_. In our example, we use a normal approximation for the sum of uniform random variables to link Δ_*t*_, and *n*. Specifically, we assume:

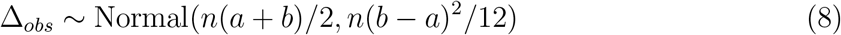

where Δ_*obs*_ is the observed gap length and *n* is the latent number of missed signals during Δ_*obs*_. In this example, *n* is an estimated parameter, while *a, b*, and Δ_*obs*_ are given as data. This approach greatly improved MCMC efficiency and provided reasonable estimates for our study system. However, a variety of approaches are possible depending on the study system. For example, the signal rate sub-model for *n* and Δ_*t*_ could be formulated as a Poisson process where *n* ~ Poisson(λ) so that Δ_*t*_ ~ Exponential(λ). This Poisson formulation may be particularly advantageous for studies focused on animal calls or cues such as the monitoring of whales or, in terrestrial systems, birds or primates. Changes in *n*, however, also cause dimensionality changes in the latent trajectory **u** as *n* is equivalent to the number of locations where a signal was emitted. As such, trajectories for all possible values of *n* are monitored and updated during each MCMC iteration. A Metropolis-Hastings update is used to accept or reject a proposed *n* conditional on its associated trajectory and other model parameters. For example, larger values of *n* are likely to be rejected when an individual is towards the interior of the array and detection probability is high. Conversely, in areas of low detection probability, *n* is primarily influenced by prior information on signal rate and duration of the observed interval (here, *a, b*, the Normal approximation equation, and Δ_*obs*_). Numerous statistical and ecological extensions to this general signal rate sub-model are possible, however, this relatively simple example demonstrates the concept of a sub-model for signal rate. Inclusion of the signal rate sub-model is not possible in JAGS (that we are aware of) and instead we developed a custom MCMC algorithm in R (Appendix A).

### 2.4 Illustration: Simulated data

We simulated 100 data sets for a system that involved 25 acoustically tagged individuals, subjected to sampling at 100 sensors on a 10 x 10 grid with unit spacing (Fig. 1). The R code for simulating this population and sampling array, as well as for fitting all the models and post-processing the output, are given in Appendix A. The initial location of each tagged individual was distributed randomly over the area in the vicinity of the array as shown in Fig. 1. Each signal interval was generated as a uniform random variable where *a* = 1 and *b* = 2 time units (e.g., minutes) and sampling occurred across 150 time units (therefore the number of potential signals of each individual is a random outcome). Obviously the time scale here is arbitrary and the system can be rescaled by increasing or decreasing the time interval. We simulated detections using the half normal model with *p*_0_ = logit^−1^ (0.25) = 0.562 and detection radius parameter to be *σ_det_* = 0.75 in the standardized units shown in Fig. 1. The standard deviation of the Brownian motion movement process, *σ_u_*, was set at 0.25. We selected these particular parameter settings so that the probability of detecting an individual within the array was > 0.90, but quickly decreased as distance from the array increased (Fig. 2). As such, individual locations near the center of the array were detected at 0 - 6 sensors, while locations on the periphery were detected at 0 - 2 sensors (Fig. 2). The true movement trajectories and detection locations of two individuals are shown in Fig. 2. We see that signals produced at interior locations rarely go undetected, individuals at interior locations are often detected at > 1 sensor, and the number of detections decreases as individuals move towards the periphery of the array. This situation illustrates one of the important motivations for using an explicit movement model in localization: the *observed* detections are distinctly biased toward the interior of the sensor array where sampling is more intensive. Therefore, localizations using classical methods will also necessarily be biased toward areas of higher sampling intensity. This sampling bias must be accounted for in studies of movement and resource selection unless sensor placement itself is random with respect to habitat structure. In an SCR framework, however, signals that produce zero detections provide information on the detection process but ignoring all-zero occasions may bias parameter estimates [18].

**Figure 1:**
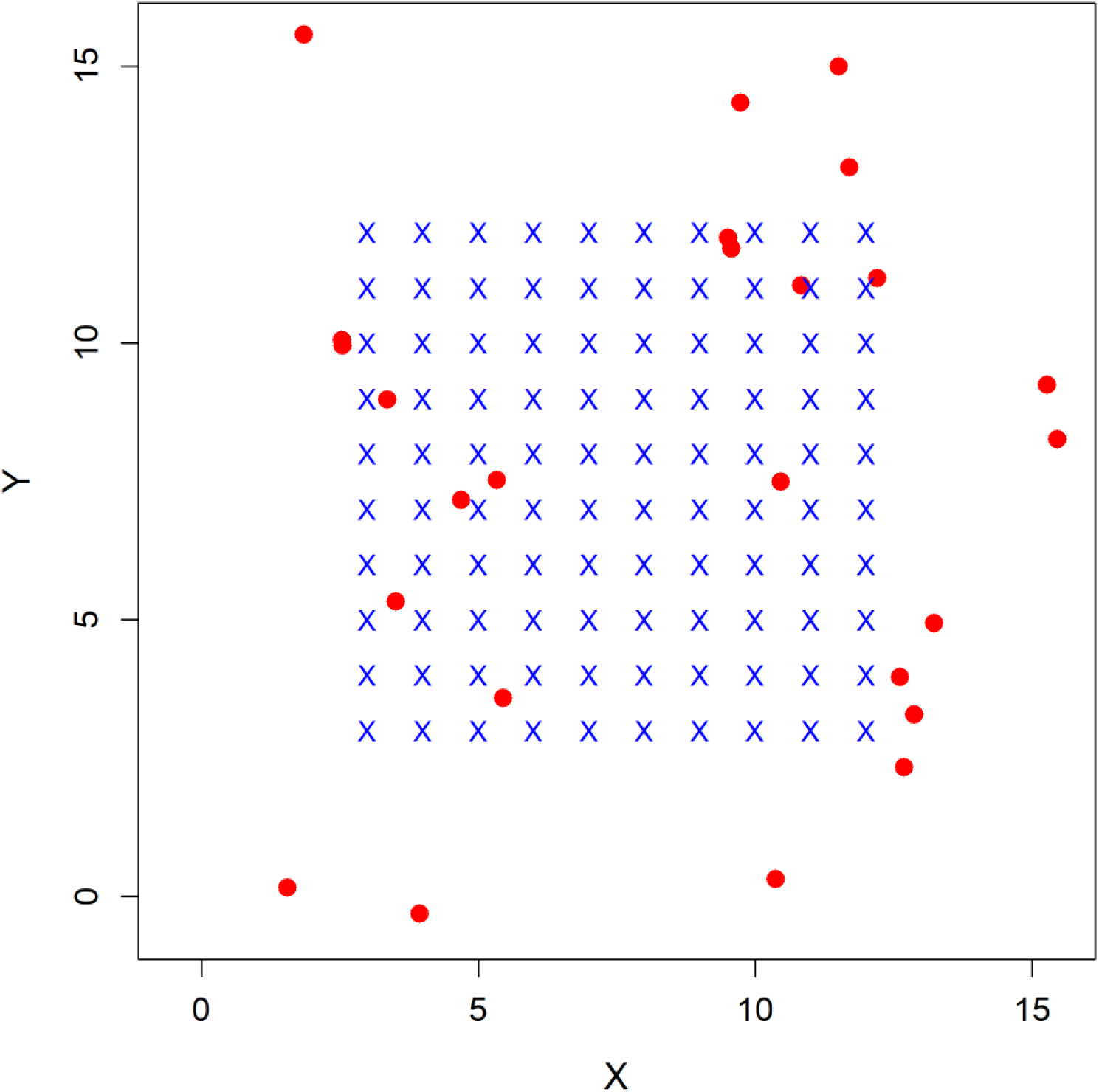
Experimental array used for simulated encounter data. Sensor array (blue x’s) has unit spacing. Initial positions (red dots) of 25 tagged individuals were simulated randomly on the plot region defined by buffering the sensor array by 5*σ_det_* units (here 3.75 units). Subsequent positions of each individual were simulated by Brownian motion as described in the main text.

**Figure 2:**
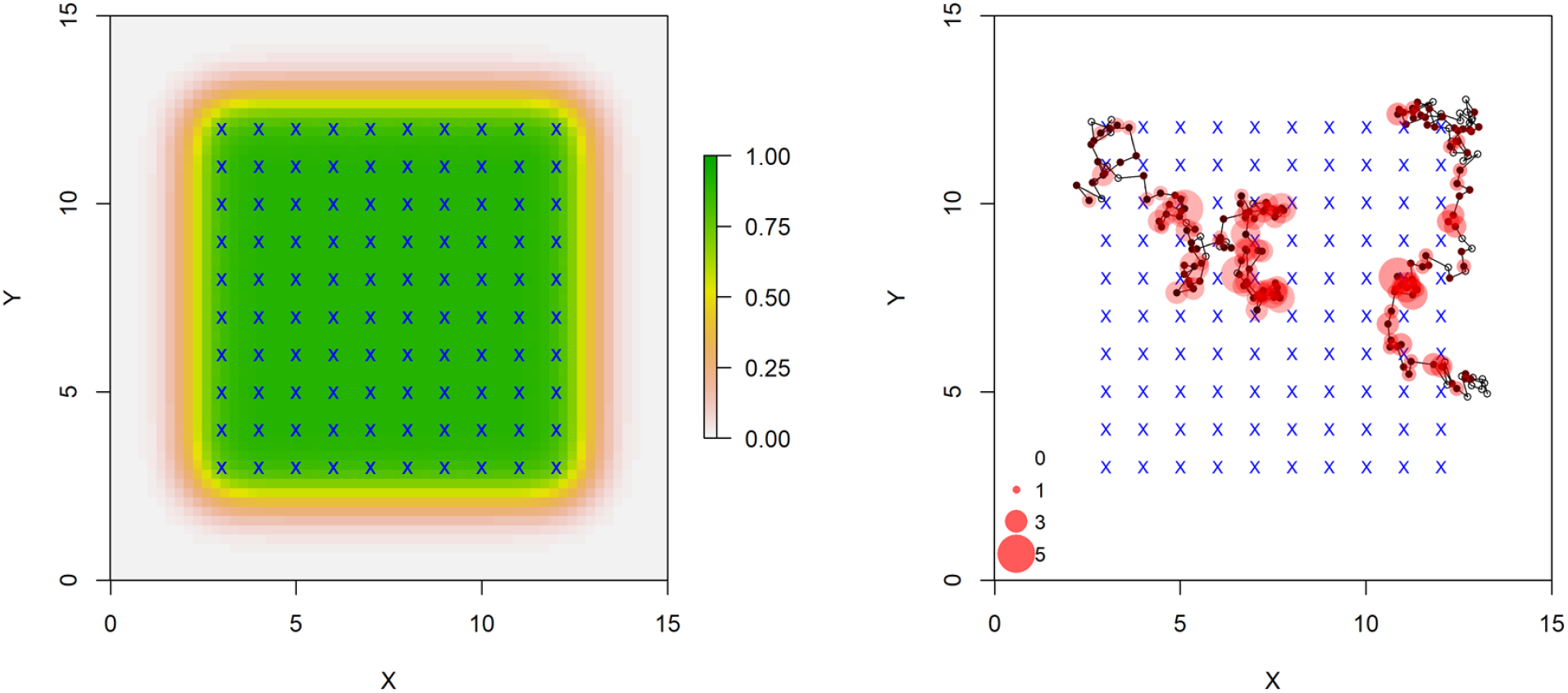
Location specific detection probability (probability of detecting a signal at ≥ 1 sensor) under current simulation settings (*σ_det_* = 0.75, *p*_0_ = 0.56; left). Sensor locations are denoted by blue x’s. True trajectories and encounter histories of two tagged individuals (right) where black dots denote locations when the acoustic tag signaled (filled [detected], open [not detected]). Locations with detections are shown in red, scaled to the number of detections (0 - 5 detections per location). The parameters *σ_det_* and *p*_0_ are easily varied to affect the expected probability of detection and the rate at which detection probability declines with distance from a sensor.

We analyzed each simulated dataset using four modelling approaches, one approach that used independent localization based on detected locations, and three types of movement-assisted localization models. The independent localization model did not involve the movement model and used only occasions with > 0 detections. Second, we fit a movement-assisted localization model using only detection occasions (hereafter movement-assisted localization detection-only model). Third, we fit a movement-assisted localization model where we assumed all interval durations (Δ_*t*_) were known even when the tag was not detected (hereafter movement-assisted localization known-interval model). Although this approach allows estimation of all undetected signal locations, we buffered the first and last detection occasions by five intervals to prevent an inordinate number of estimated leading and trailing locations. Finally, we fit a movement-assisted localization model where the number of missed signals and their associated intervals are unknown and are instead estimated as part of the model (hereafter movement-assisted localization unknown-interval model). While this movement-assisted localization unknown-interval model is more complex, it likely describes the most realistic field sampling protocol. We compared these four modeling approaches to investigate biases in the SCR framework, improvements in localization due to the integration of a movement model, effects of ignoring all-zero occasions, and technological (or statistical) considerations if we can assume the number of unobserved signals is known or partially informed by transmitter settings.

We fit the first three models using the software JAGS [35] accessed through R version 3.5.2 [36], using the jagsUI package [37], with specifications similar to those shown in Panel 1 (see Appendix A for the full R script). We ran three parallel Markov chains for 15 000 iterations with 5000 burn-in iterations and 2000 adaptation iterations. Chains were thinned by 2 to reduce the size of model output. The movement-assisted localization unknown-interval model required a custom MCMC algorithm implemented in R (see Appendix A for the full R script) due to the challenges of the signal rate sub-model for updating the latent number of missed signals and their associated intervals and locations. Due to slower mixing, the movement-assisted localization unknown-interval model used three chains of 10 000 burn-in iterations and 50 000 saved iterations thinned by 10. We summarize posterior distributions as medians and 2.5 and 97.5 percentiles (95% CRI). We evaluated relative frequentist bias, using posterior means as point estimates, for higher level parameters, (*σ_det_*, *σ_u_*, *p*_0_). We expect some bias in the independent localization model and detection-only model as these methods discard the all-zero encounter occasions. To characterize localization efficiency, we evaluated three metrics describing the accuracy and precision of (i) mean posterior location estimates relative to the true locations, (ii) precision of the posterior localizations, and (iii) localization credible interval coverage. To evaluate precision of means, we computed the Euclidian distance between each posterior mean and the true location (RMSE). For precision of the full posteriors, we calculated the Euclidian distance between each posterior estimate and the true location (hereafter precision). In both cases, smaller values are preferred. We also recorded the proportion of occasions in which the true location was within the 95% highest posterior density kernel, which we report as localization coverage.

## 3 Results

### 3.1 Simulation results

Both the movement-assisted localization known-interval and unknown-interval models returned relatively unbiased estimates of the movement and detection parameters (Table 1). The independent localization and movement-assisted localization detection-only models displayed positive bias in p0 due to the exclusion of all-zero occasions (Table 1). In the detection-only model, *σ_det_* and *σ_u_* displayed −1% and −9% relative bias, respectively (Table 1). The unknown-interval model reduced the relative bias in *σ_det_* and *σ_u_* to 0% and −8.0%, respectively (Table 1), while reducing bias in *p*_0_ from 16% to −1% and providing additional inference to locations with zero detections.

**Table 1:** Mean and relative bias of movement (*σ_u_*) and detection (*σ_det_*, *p*_0_) parameters from 100 simulated data sets analyzed using the independent localization model (Ind. loc) and three forms of movement-assisted localization using: (Mvmt DO) detection occasions only, (Mvmt KI) known signal intervals, or (Mvmt UI) unknown signal intervals, which are modeled as random variables.

| Param. | True | Ind. loc |  | Mvmt DO |  | Mvmt KI |  | Mvmt UI |  |
| --- | --- | --- | --- | --- | --- | --- | --- | --- | --- |
|  |  | Mean | Bias | Mean | Bias | Mean | Bias | Mean | Bias |
| $\sigma_u$ | 0.25 | NA | NA | 0.23 | -0.09 | 0.25 | 0.00 | 0.23 | -0.08 |
| $\sigma_{det}$ | 0.75 | 0.79 | 0.06 | 0.74 | -0.01 | 0.75 | 0.00 | 0.75 | 0.00 |
| $p_0$ | 0.56 | 0.62 | 0.11 | 0.65 | 0.16 | 0.56 | -0.01 | 0.56 | -0.01 |

We observed large benefits in our primary objective of improving localization by integrating an underlying movement model. First, the three movement-assisted localization modeling approaches reduced RMSE by nearly one-half relative to the independent localization model when the number of detections were < 3, and still provided noticeable improvements when the number of detections was > 3 (Table 2, Fig. 3). Movement-assisted localization models achieved similar levels of RMSE and precision using 1 - 2 detections as the independent localization model achieved with 5 - 6 detections (Table 2). Credible interval coverage of ut was generally > 0.90 for all models across all levels of detections (1 - 6 detections per occasion; Table 2).

**Figure 3:**
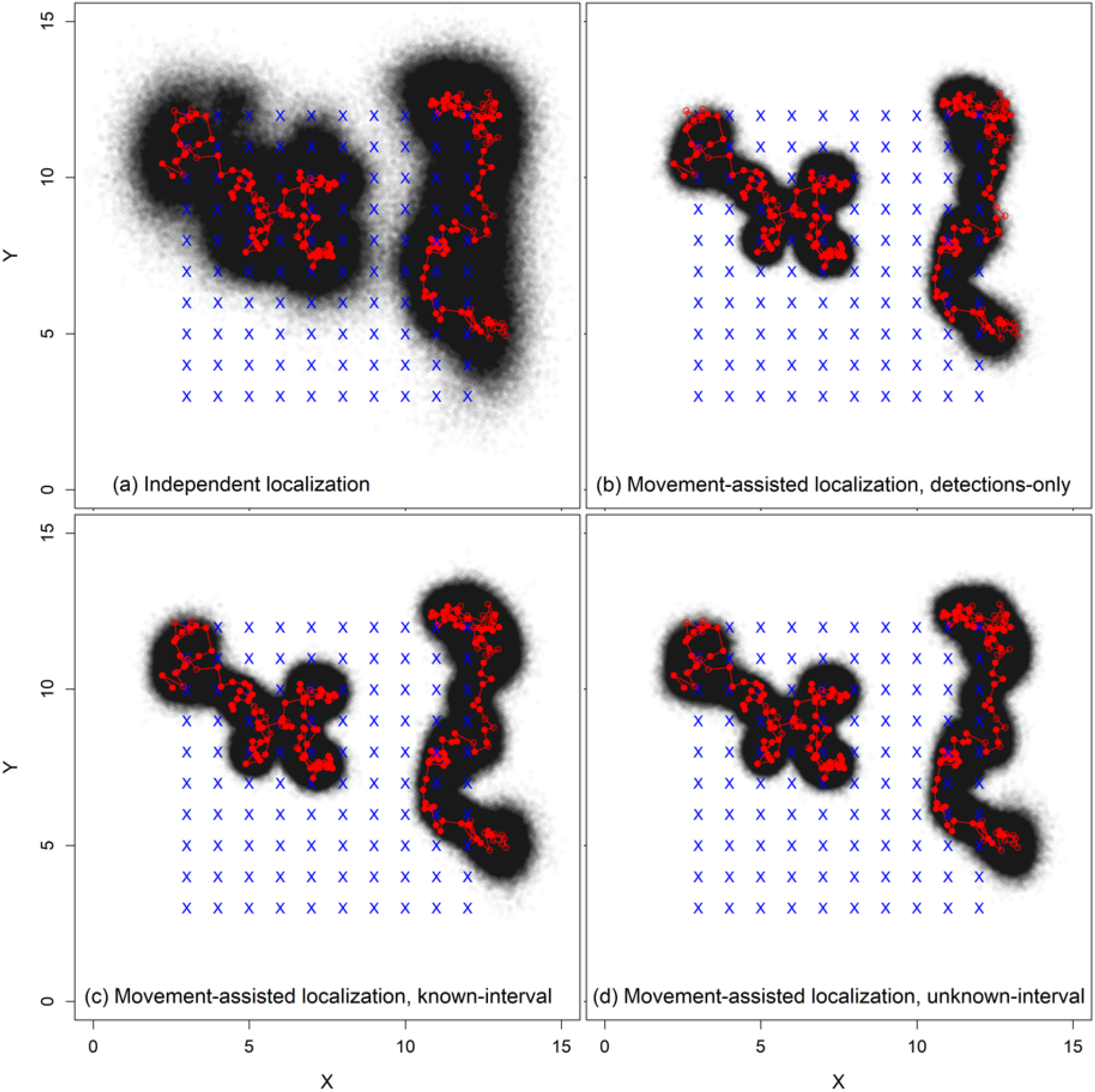
Latent trajectories of two individuals (red) with detection and non-detection locations (filled and open red dots, respectively; see Fig. 2 for additional details) and modelspecific posterior localizations (2000 posterior samples; black points). Modeling approaches include (a) the independent localization model and (b - d) three forms of movement-assisted localization using: (b) detection occasions only, (c) assuming all time intervals are known, or (d) modeling the unknown time intervals as random variables. Sensor array is denoted by blue x’s.

**Table 2:** Root mean squared error of posterior means (RMSE), precision of full posterior (Prec; smaller is better), and credible interval coverage (Cov) for localization of ut from 100 simulated data sets analyzed using the independent localization model (Ind loc) and three forms of movement-assisted localization: (Mvmt DO) detection occasions only, (Mvmt KI) assuming the signal interval is known, or (Mvmt UI) modeling the unknown signal interval as a random variable. Values are presented as a function of the number of sensors where the signal was detected. Only the known-interval model localizes to a specific ut when 0 detections. Due to their rarity, results for signals detected at 7 or 8 sensors were excluded from results.

| Det | Ind loc |  |  | Mvmt DO |  |  | Mvmt KI |  |  | Mvmt UI |  |  |
| --- | --- | --- | --- | --- | --- | --- | --- | --- | --- | --- | --- | --- |
|  | RMSE | Prec | Cov | RSME | Prec | Cov | RMSE | Prec | Cov | RMSE | Prec | Cov |
| 0 | NA | NA | NA | NA | NA | NA | 0.72 | 1.00 | 0.94 | NA | NA | NA |
| 1 | 0.89 | 1.30 | 0.96 | 0.49 | 0.63 | 0.88 | 0.42 | 0.60 | 0.95 | 0.55 | 0.74 | 0.92 |
| 2 | 0.63 | 0.91 | 0.96 | 0.37 | 0.50 | 0.92 | 0.36 | 0.51 | 0.95 | 0.46 | 0.63 | 0.93 |
| 3 | 0.51 | 0.74 | 0.96 | 0.33 | 0.45 | 0.93 | 0.33 | 0.46 | 0.95 | 0.42 | 0.58 | 0.94 |
| 4 | 0.45 | 0.65 | 0.96 | 0.30 | 0.42 | 0.93 | 0.31 | 0.43 | 0.95 | 0.39 | 0.54 | 0.94 |
| 5 | 0.40 | 0.58 | 0.96 | 0.29 | 0.40 | 0.94 | 0.29 | 0.41 | 0.95 | 0.38 | 0.52 | 0.92 |
| 6 | 0.37 | 0.53 | 0.96 | 0.26 | 0.37 | 0.93 | 0.26 | 0.38 | 0.93 | 0.36 | 0.49 | 0.94 |

Inference to locations with zero detections is possible in both the movement-assisted localization known-interval and unknown-interval models. The known-interval model returned relatively precise estimates of zero-detection locations and achieved close to nominal credible interval coverage (0.94, Table 2). Similarly, the unknown-interval model accurately estimated the number of missed signals per individual (relative bias = 2%) across a wide range of values (< 5 to > 100 missed signals per individual; Fig. 4). The unknown-interval model provides a set of posterior locations conditional on the latent number of missed signals (rather than a fixed number of locations as in the known-interval model), and as such RMSE and precision for specific zero-detection locations are not directly comparable (Table 2). Minimally biased estimates and close to nominal credible interval coverage of upper level parameters, missed signals, and localization (Table 1, Table 2, Fig. 4) together demonstrate the capacity of the unknown-interval model to simultaneously estimate movement, signal rate, and detection processes from acoustic data even when the numbers of missed signals are unknown.

**Figure 4:**
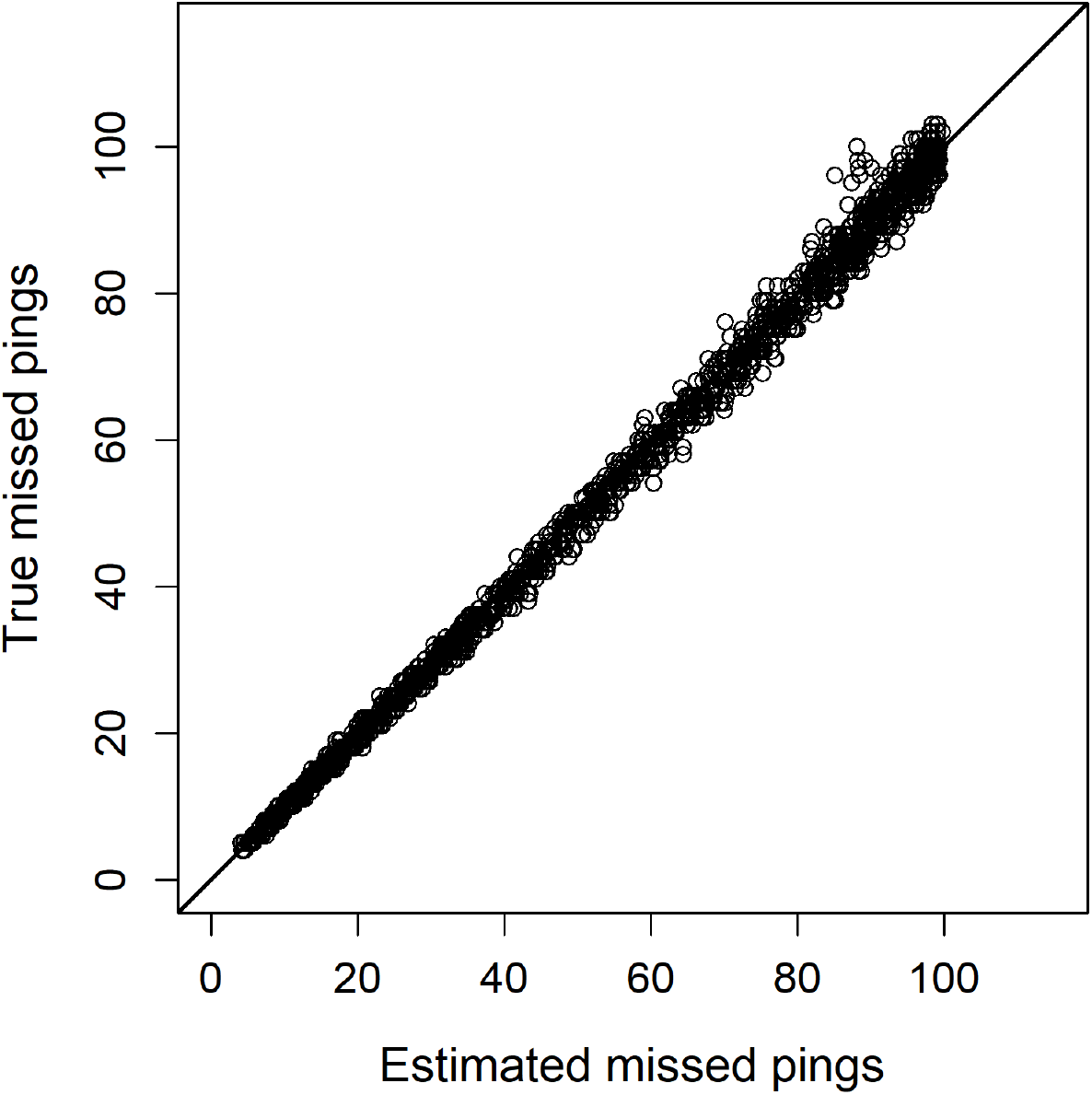
Number of true missed signals per simulated individual (100 simulated data sets of 25 individuals; y-axis) and the number of estimated missed signals from the movement-assisted localization unknown interval model (x-axis). Diagonal line is a 1:1 line. Relative bias = 2%.

Integrating the movement process dramatically improved precision of posterior localizations while still maintaining nominal or close to nominal credible interval coverage (Table 2, Fig. 3). Increased localization precision between the independent localization model and the movement-assisted localization detection-only model are due directly to the integration of the movement model (Fig. 3 a vs b). The known-interval model further extends inference to locations with zero detections, thus the full posterior trajectory is slightly larger than the detection-only model (Fig. 3 c vs b). Finally, the unknown-interval model relaxes the requirement of a known number of missed signals and the interval between those signals (Fig. 3 d). Interestingly, localization at the level of the full trajectory was only minimally influenced by an unknown signal interval in these examples (Fig. 3 c vs d).

The primary influence of the movement model is the restriction of locations to regions consistent with the movement trajectory and preventing unrealistically large movements between detection occasions, which results in improved accuracy and precision of the full trajectory (Fig. 3). An animation of occasion-specific localizations are provided in Appendix B, further demonstrating the improved localization from movement-assisted localization methods at the single-occasion and full trajectory levels. For the unknown-interval model, animations present the marginal distribution of ut when intervals consist of multiple zero detection occasions.

## 4 Discussion

We described a localization approach for acoustic telemetry data that combines classical ideas of movement modeling [27] and localization of sources from sensor arrays. Similar ideas exist in the field of spatial capture-recapture [18, 38]. In fact, localization of sources in acoustic telemetry is precisely analogous to estimation of an individual’s activity center in terrestrial spatial capture-recapture and the movement of activity centers through time [18, 21, 25, 26]. The relevance of the movement model, however, is of much greater importance in acoustic telemetry studies where a high frequency of observations can introduce considerably more information about individual locations over short time periods. Conversely, in many terrestrial capture-recapture studies spatial sampling is much coarser and temporal observations usually occur at a much lower frequency, thus the movement process itself often provides less information about individual locations and model parameters [21].

Localization from sensor array data only requires the spatial encounter history as we have demonstrated in this paper. However, it is possible to use auxiliary information on signal strength [22], time-difference-of-arrival (TDOA), or even both signal strength and TDOA simultaneously [15, 20]. Such auxiliary information often improves the precision of localizations, and can be included in the localization model as independent data with contribution to the likelihood depending on the nature of the auxiliary data [17, 20]. For example, TDOA is incorporated into the localization model by regarding arrival time of the *t^th^* signal at detector j as a normal random variable with mean *E*(*x_t,k_*|**u**_*t*_) = *β*_0*t*_ + *β*_1_||**x** − **u**_*t*_|| or some similar function of distance, and normal measurement error of arrival time with variance 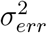. Here, *β*_0*t*_ is the time the *t*th sound was generated and *β*_1_ is the inverse of the speed of sound in water or air depending on the context. Because *β*_0*t*_ is not known, the likelihood can be expressed in terms of the time *difference* between arrival at *j* and the mean arrival time. This requires at least 2 measurements of arrival time are obtained for each localization [20]. Any model for localization based on auxiliary data, however, can be integrated into our movement-assisted localization framework.

Classical localization based on hyperbolic positioning algorithms requires that individuals are detected simultaneously at multiple sensors, and improvements in localization precision are primarily achieved through increased density of sensors. Alternatively, we described a method that uses information on previous and subsequent locations of an individual to produce improved localizations with fewer detections, including even zero detections. Conceptually, our approach to movement-assisted localization is based on formulating the model for an individual’s spatial-temporal detection history conditional on the individual’s latent movement trajectory (i.e. its sequence of locations). This formulation is similar to specification of the likelihood in spatial capture-recapture models and we feel that it has a number of benefits to modeling acoustic telemetry data, including joint inference about the entire trajectory of an individual and model parameters based on all available data. Additionally, this approach allows for *in situ* estimation of the detection range parameter(s) in the model simultaneously with estimation of the latent trajectory. The framework can be extended to allow heterogeneity in detection range due to environment and other factors. For example, detection probability can be expressed as a function of covariates such as depth or distance from shore, or non-Euclidean effective distance models which allow for sound attenuation in heterogeneous environments [32].

A second important application of our model is in developing improved models of resource selection, habitat use, and identification of movement or migration corridors. Resource selection models usually regard individual locations as fixed known points, which is a reasonable assumption in classical terrestrial GPS-based telemetry [although see 39]. However, in acoustic telemetry applications locations of individuals are unobserved and inferred based on detections at known sensor locations. Considerable localization uncertainty reduces the power to detect important habitat effects. Our approach estimates the posterior distribution of the individual’s trajectory (Fig. 3), which yields a characterization of the latent point pattern of *true* locations of the individual during the study. The underlying point pattern of true locations can then be used to obtain a posterior characterization of any particular resource selection model conditional on the point locations. Thus, a coherent accounting for uncertainty in locations can propagate through to the inference about resource selection and movement processes, providing a solution to explicitly accommodating the variable precision of localizations in models of resource selection and similar applications. Moreover, our movement-assisted localization approach accommodates inherent bias in acoustic localization data that is due to estimated locations being biased toward areas of high sensor density (e.g., toward the interior of an array vs. the edge).

A third potential application of this framework is for modeling the number of active tags in the vicinity of the array (i.e. abundance). In practice, the number of tagged individuals available for capture is rarely known. This is precisely the problem that spatial capture-recapture resolves by introducing parameter(s) to describe the density of the underlying point process and a key motivating factor for viewing the problem as one of SCR. Here, SCR provides a framework to localize not only to detected transmitters, but also estimate the density and distribution of undetected transmitters. Further, it may then be possible to explicitly model the effect of density dependence on detection probability. When multiple tags of the same frequency exist in the vicinity of a sensor array, interference can induce a density dependent decrease in detection probability. We believe this is an important research area which can be resolved using our formulation of the movement-assisted localization model.

SCR methods provide a hierarchical framework to describe the three primary components of acoustic telemetry: (1) the movement of individuals, (2) the signal or cue rate, and (3) the detection process. Our simulation study described a relatively simple example to demonstrate SCR as a method to analyze acoustic telemetry data, and by no means explores all the possible applications or challenges. The greatest challenges to these methods arise from an unknown number of missed signals and the development of an appropriate signal rate sub-model. In many acoustic telemetry systems, the inter-signal duration is defined by transmitter settings and can encompass large periods of time. Under these settings, model complexity quickly increases due to the high number of possibly missed signals (*n*), the inter-signal intervals (Δ_*t*_), and associated changing dimensions of trajectory locations (**u**_*t*_). Broadly speaking, this is not a deficiency in the application of SCR methods but a restriction on the specific model described herein. For example, one approach to situations where tracking specific Δ_*t*_ are difficult or even unnecessary for inference, is to standardize Δ_*t*_ to a fixed interval (e.g., 10-minute or 1-hr periods) and model localizations as the average location during an interval [40]. Overall, most of these challenges can be addressed by modifications to the modelling approaches we described, while retaining the general SCR framework and hierarchical approaches developed herein. Alternatively, some of these sampling challenges may be addressed by design or technological advances (e.g., by fixing the sampling interval or retaining data on the transmission sequence). In most situations, resolving uncertainties through careful planning and design is highly preferable compared to trying to resolve uncertainties with statistical models.

Our application of movement-assisted localization focused on acoustic telemetry data, however, similar concepts are applicable to acoustic monitoring systems aimed at detecting signals, calls, or cues. In acoustic monitoring applications signal intervals are stochastic, being equivalent to the signal rate of individuals in the population. It is therefore essential to develop models for cue rate in order to make progress toward developing unbiased abundance and demographic parameter estimates from acoustic data. Our unknown-interval model does just this, as it estimates the number of signals (e.g., signals emitted from a tag [active] or cues produced by an animal [passive calls, clicks, singing, etc.]) and the duration between signals as part of the model. Prior information on cue rates may arise in the form of transmitter settings or auxiliary data on cue rates. In our example, integrating the latent movement, detection, and cue rate processes allowed estimation of all parameters with minimal data. Our movement-assisted localization framework is a particularly promising approach to estimate detection probability, a key step to accurate abundance estimation in any system [41]. Further, describing these process through an underlying point process model and implementation in a Bayesian framework provides tremendous flexibility to integrate auxiliary information to improve parameter estimation and explore additional ecological processes (e.g., aerial- or boat-based count data to inform abundance, call rate data to inform cue rates, telemetry data to inform movement). Joint efforts from ecologists, statisticians, and engineers will be vital to solving these challenges and advance the utility of acoustic studies to address pressing challenges in the study of movement, survival, and abundance in both the aquatic and terrestrial realms.

Finally, our movement-assisted localization framework provides a host of opportunities to evaluate integrated, multi-objective study designs. In general, the design of sensor arrays can be based on objectives that involve maximizing the probability of detecting a tagged individual or statistical precision of detection parameters [18, 42]. Combining a state-space movement model with a sub-model for the detection process [i.e. a positioning model; 17] dramatically improved localization precision, whereby only 1 - 2 detections per occasion resulted in similar precision as 5 - 6 detections in the independent localization model. As such, study designs may consider coarser sensor spacing and broader spatial coverage without sacrificing localization precision. Overall, movement-assisted localization provides a flexible, transparent framework that is easily adapted to aquatic telemetry systems and studies interested in using acoustic data to investigate ecological processes.

## 5 Conclusions

Understanding animal movement and space-use is crucial for effective conservation and management of species. Acoustic telemetry data are increasingly used to study aquatic ecosystems and face many similar logistical and modeling challenges as spatial capture-recapture studies in terrestrial environments. Our results demonstrate a unifying framework to model acoustic telemetry data based on an underlying spatial capture-recapture model integrated with explicit movement and signal rate sub-models. This approach improves localization estimates and can be adapted to a variety of species- and study-specific settings such as unknown signal (or cue) rates, imperfect detection, and the integration of habitat data to inform individual- and population-level movement and space-use.

## Additional files

**Appendix A** R code for simulating movement, signal rate, and the detection process describe in this paper, as well as for fitting all the models and some post-processing.

**Appendix B** Animation of the latent trajectories for two individuals. Animations denote detection and non-detection locations (filled and open red dots, respectively; see Fig. 2 for additional details) and model- and occasion-specific posterior localization estimates (black points). Modeling approaches include (a) the independent localization model and (b - d) three forms of movement-assisted localization: (b) detection occasions only, (c) assuming all time intervals are known, or (d) modeling the unknown time intervals as random variables. Sensor array is denoted by blue x’s and sensors that recorded a detection at each time step are highlighted in red.

## Abbreviations

MCMC: Markov chain Monte Carlo
RMSE: Root-mean squared error
SCR: Spatial Capture-recapture
TDOA: time-difference-of-arrival

## Acknowledgments

The authors thank L. Duskey, D. Prosser, and D. Hondorp for helpful comments and suggestions on this manuscript. Any use of trade, product, or firm names is for descriptive purposes only and does not imply endorsement by the U.S. Government.

## Authors contributions

JAR initiated this study. JAR and NJH developed models and conducted data analyses. NJH and JAR wrote the manuscript. All authors approved of the final manuscript submission.

## Availability of data and materials

All data generated and analysed during this study are included in this published article [and its supplementary information files].

## Ethics approval and consent to participate

Not applicable.

## Consent for publication

Not applicable.

## Footnotes

1 In keeping with the terminology in spatial capture-recapture, we refer to this planar region as the statespace of the point process which defines the potential locations of individuals during the study. In practice, the state-space should be chosen to be much larger than the region containing the sensor array because individual locations may be detectable some distance from the sensors on the boundary.

