## Appendix B for "Movement-assisted localization from acoustic telemetry data"

### Slide 1
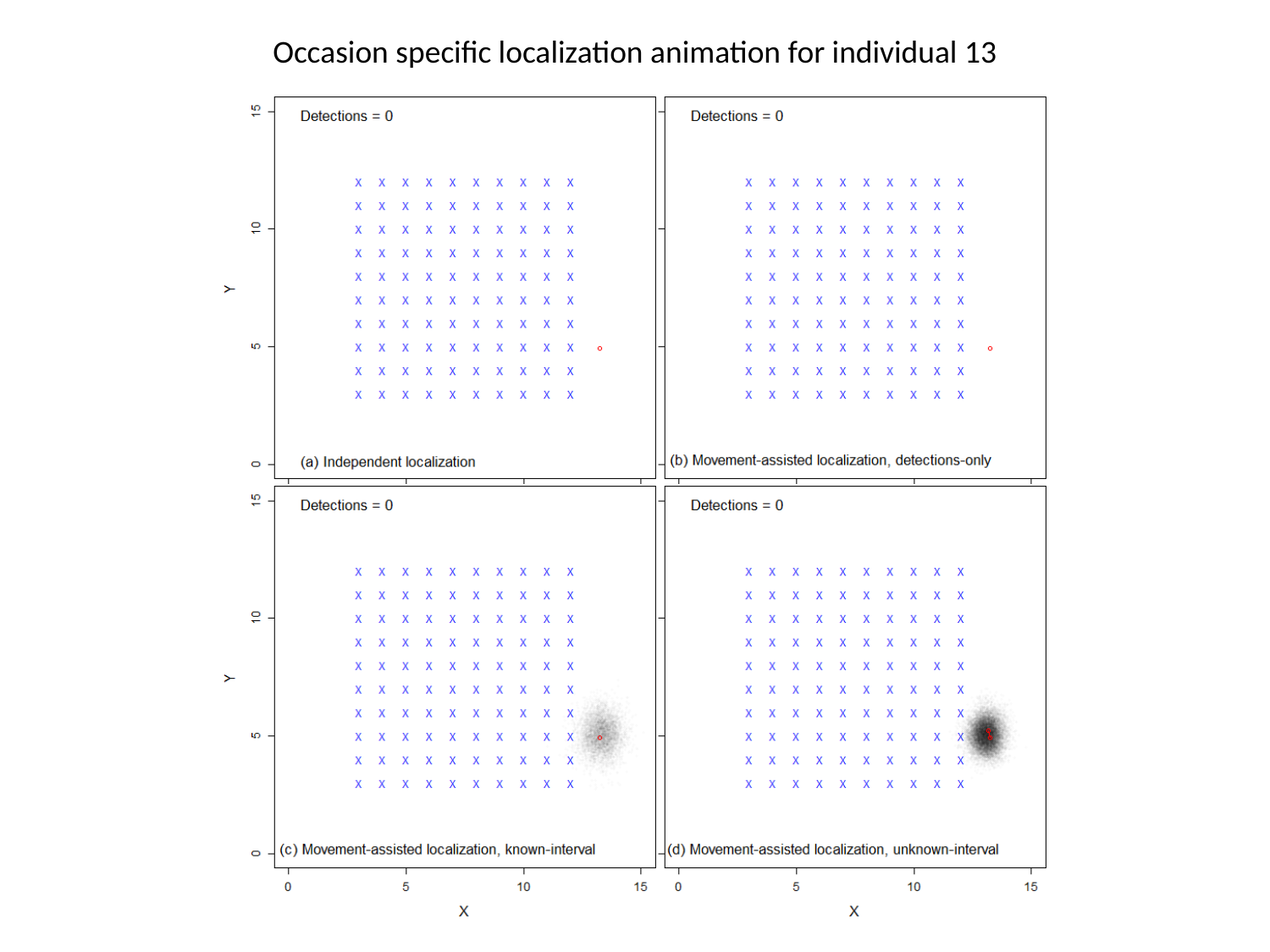

Occasion specific localization animation for individual 13

### Slide 2
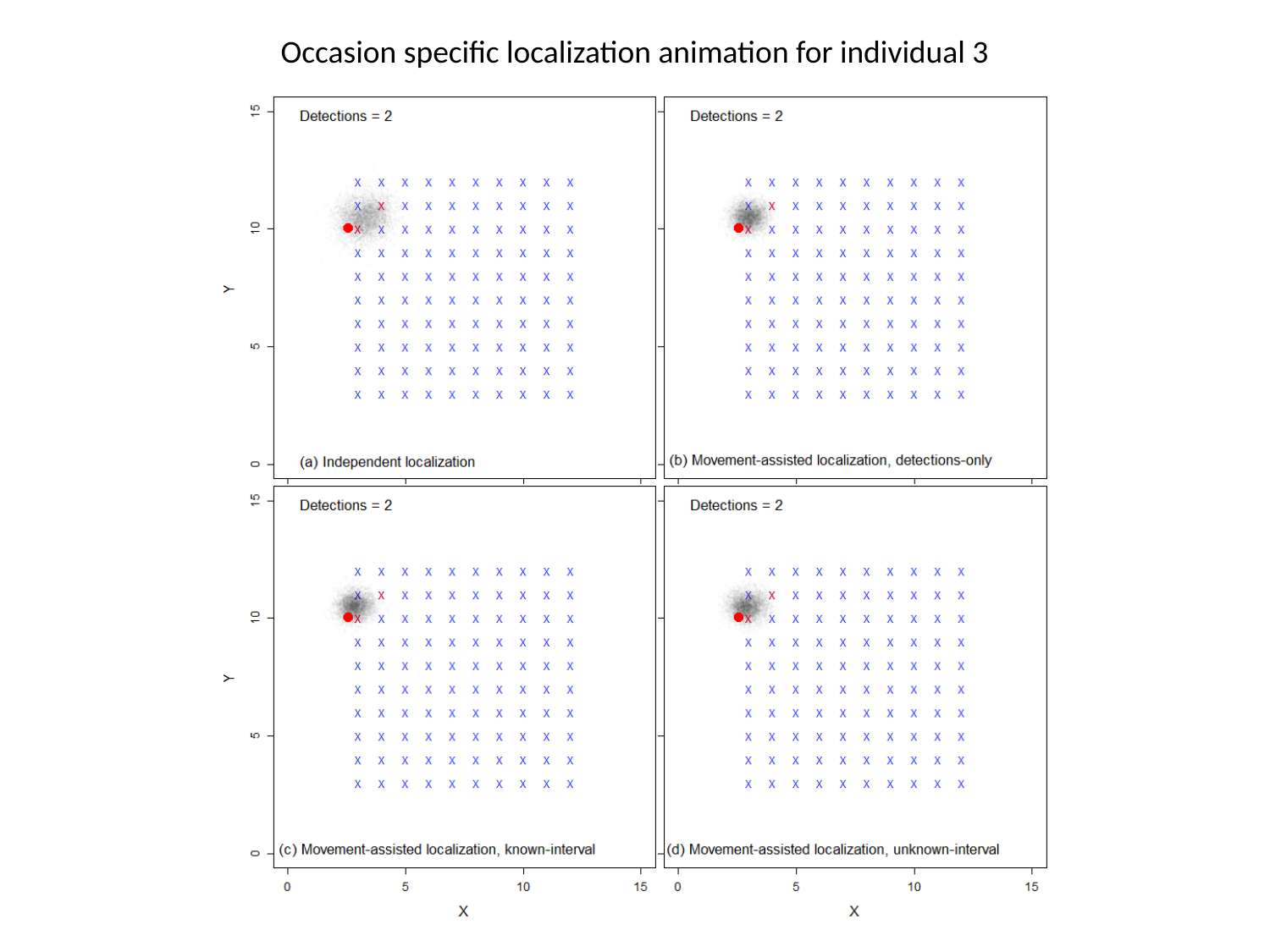

Occasion specific localization animation for individual 3
